## Supplementary Materials for "Membrane to cortex attachment determines different mechanical phenotypes in LGR5+ and LGR5- colorectal cancer cells"

### Supplementary Figures

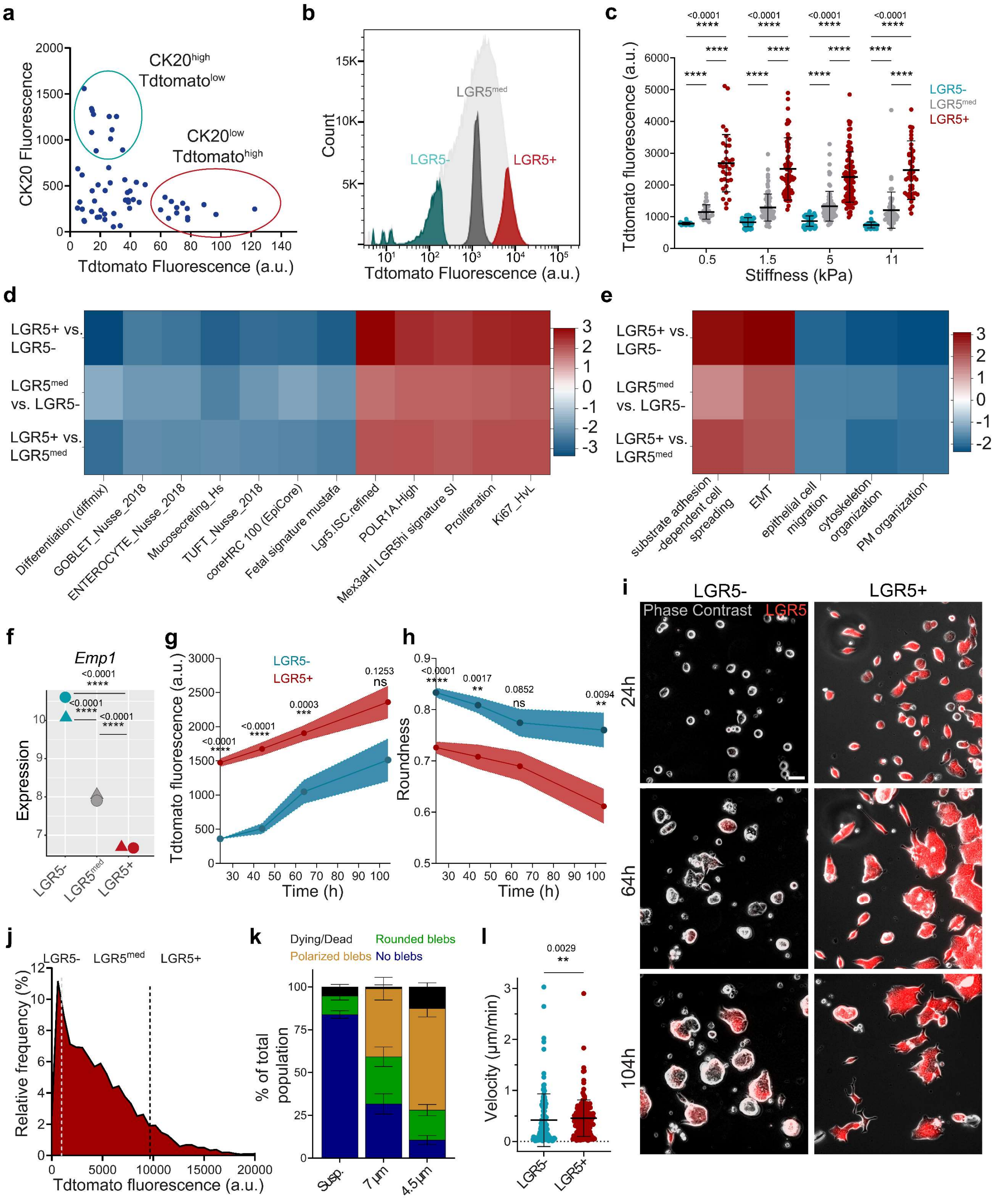

**Supplementary Figure 1. Plasticity of PDO7 single cells, LGR5-Tdtomato fluorescence and response to confinement. a.** Quantification of CK20 and Tdtomato fluorescence in CRC PDOs cultured for 1 week in culture matrix gel. **b.** Flow cytometric sorting strategy used to obtain LGR5-, LGR5$^{med}$ and LGR5+ cells from CRC PDOs cultured for 1 week in culture matrix. **c.** Mean

Tdtomato fluorescence intensity of sorted LGR5-, LGR5$^{med}$ and LGR5+ cells 24 h after seeding on collagen I coated gels. Quantification from confocal images. Data are represented as the mean ± s.d. of n > 73 cells/condition from four independent experiments. Statistical significance was determined using Shapiro-Wilk normality test, followed by a Kruskal-Wallis multiple-comparison test. **d.** Heatmap showing NES for selected gene sets in LGR5-, LGR5$^{med}$ and LGR5+ cells. Comparison of signature scores between samples was assessed using t-tests. *p* values are listed in Table 1. **e.** Heatmap showing NES for selected GOBP gene sets in LGR5-, LGR5$^{med}$ and LGR5+ cells. Comparison of signature scores between samples was assessed using t-tests. *p* values are listed in Table 1. **f.** *Emp1* normalized expression in LGR5-, LGR5$^{med}$ and LGR5+ cells as quantified by Bulk RNA-seq. Statistical analysis was performed using t-tests. (**g, h**) Change in Tdtomato fluorescence and roundness in sorted single cells as a function of time. For each time point n ≥ 86 cells. Statistical significance was determined using two-way analysis of variance, followed by a Šidák multiple-comparison test. **i.** Time lapse of LGR5- and LGR5+ sorted single cells on 3kPa gels coated with Collagen I. Acquisition started 24 h after sorting (time 0). Representative images from two independent experiments. **j.** Tdtomato fluorescence intensity of LGR5-, LGR5$^{med}$ and LGR5+ cells in RT-DC. **k.**Percentage of dying/dead, polarized blebs, rounded blebs and no blebs in LGR5- and LGR5+ cells in suspension or under 7 and 4.5 µm confinement on a non-adhesive surface. Data are represented as the mean ± s.d. of n > 30 positions/condition. Statistical significance was determined using two-way analysis of variance, followed by a Šidák multiple-comparison test. **l**. Migration speed of tracked nuclei of LGR5- and LGR5+ cells. Data are represented as the mean ± s.d. of n > 110 cells/condition from four independent experiments. Statistical significance was determined using Shapiro-Wilk normality test, followed by a Kruskal-Wallis multiple-comparison test.

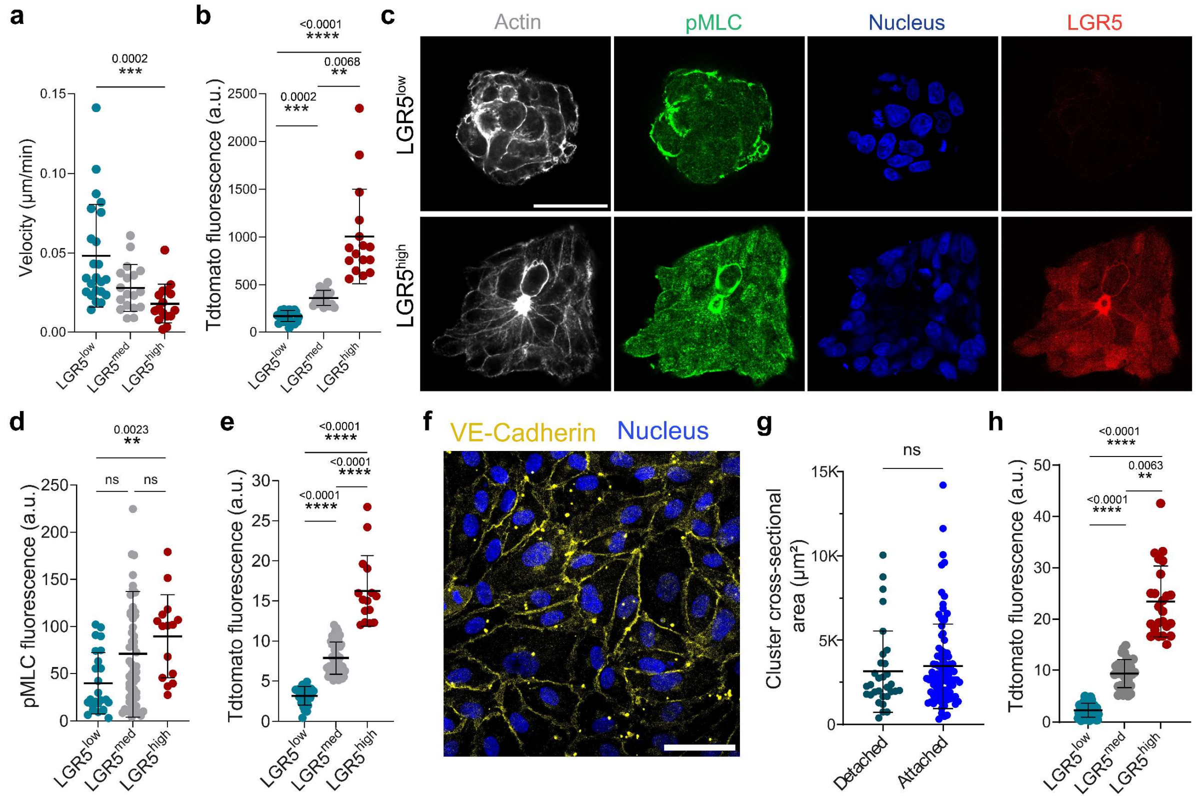

**Supplementary Figure 2. Analysis of cluster dynamics.** (**a, b**). Migration velocity (**a**) and Tdtomato fluorescence intensity (**b**) of clusters on 11kPa gels. Data are represented as the mean ± s.d. of n = 58 clusters from two independent experiments. Statistical significance was determined using Shapiro-Wilk normality test, followed by a Kruskal-Wallis multiple-comparison test. **c.** PDO clusters seeded on 3kPa gels coated with Collagen I. Clusters were stained for Actin, phosphorylated myosin light chain (pMLC) and nuclei (hoechst). LGR5+ cells are labelled with Tdtomato. Scale bar, 50 µm. (**d, e**) Quantification of mean pMLC (**d**) and Tdtomato fluorescence intensity (**e**) of $LGR5^{low}$, $LGR5^{med}$ and $LGR5^{high}$ clusters. Data are represented as the mean ± s.d. of n = 98 clusters from three independent experiments. Statistical significance was determined using Shapiro-Wilk normality test, followed by a Kruskal-Wallis multiple-comparison test. **f.** HUVEC monolayer grown for 4 days on collagen I coated gels and stained for VE-cadherin and nuclei (hoechst). Scale bar 50 µm. **g** Cross-sectional area of clusters that remained attached or detached from endothelial monolayer during the 15 h timelapse acquisition. **h.** Tdtomato fluorescence intensity of clusters divided into three groups. (**g, h**) Data are represented as the mean ± s.d. of n > 110 clusters from four independent experiments. Statistical significance was determined using Shapiro-Wilk normality test, followed by a Kruskal-Wallis multiple-comparison test.

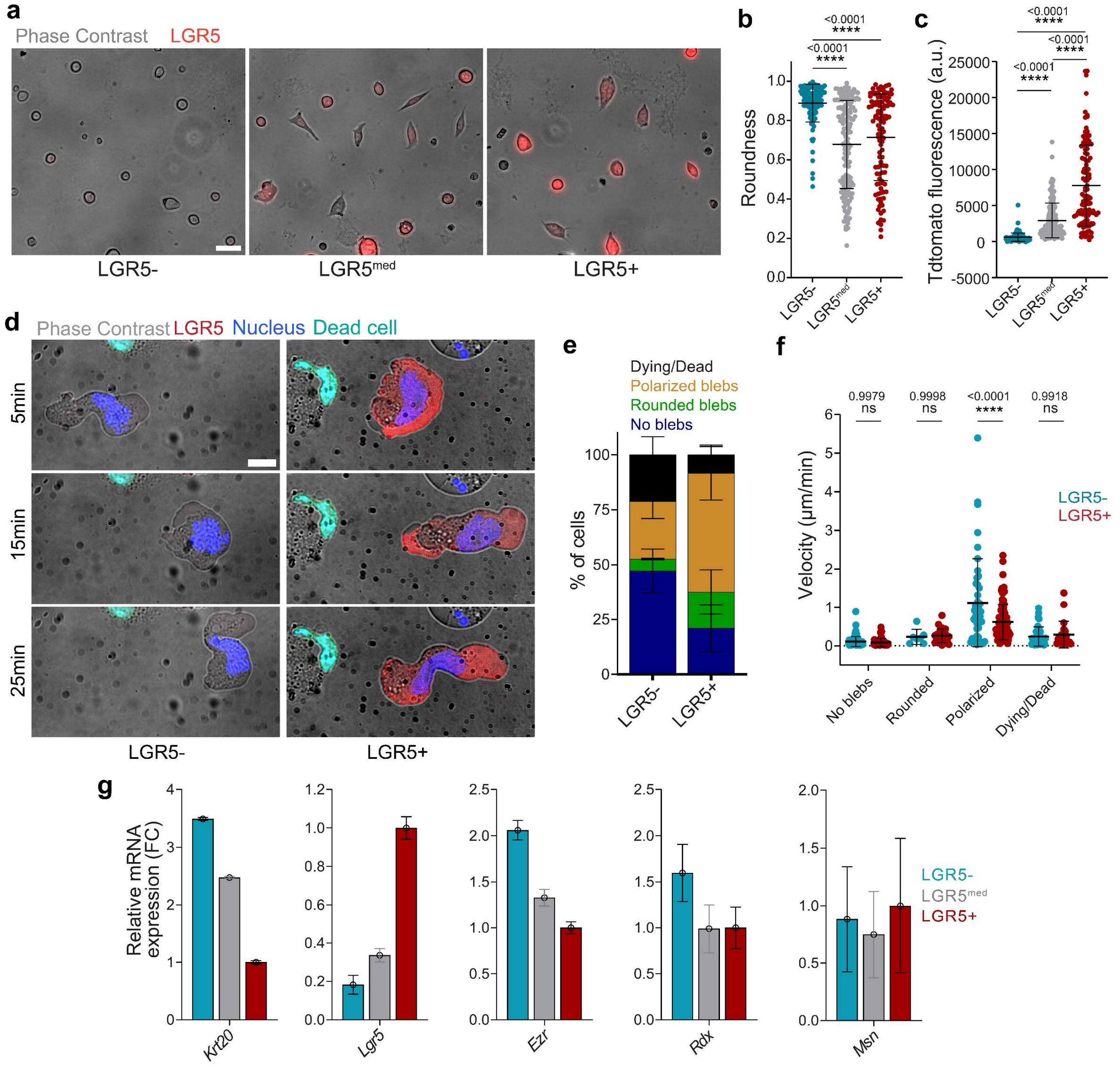

**Supplementary Figure 3. PDO-p18 cells display mechanical phenotypes similar to PDO7 cells. a.** LGR5-, LGR5$^{med}$ and LGR5+ PDO-p18 cells on 3 kPa gel substrates. Representative images of 2 independent experiments. Scale bar, 20 µm. **b.** Cell roundness measured for sorted PDO-p18 single cells seeded on collagen-I coated gel substrates of 3 kPa in stiffness. **c.** TdTomato Fluorescence of LGR5-, LGR5$^{med}$ and LGR5+ PDO-p18 cells. (**b, c**) Data are represented as the mean ± s.d. of n ≥113 cells/condition from two independent experiments. Statistical significance was determined using Shapiro-Wilk normality test, followed by a Kruskal-Wallis multiple-comparison test. **d.** Representative time lapse images of LGR5- no blebs, LGR5+ and LGR5- polarized blebs. Scale bar, 10 µm. In (**f**) and (**h**) images are representative of four independent experiments, with a total of 359 cells. **e.** Percentage of dying/dead, polarized blebs, rounded blebs and no blebs in LGR5- and LGR5+ cells under 4.5 µm confinement on a non-adhesive surface. Data are represented as the mean ± s.d. of percentages from four independent experiments. Statistical significance was determined using two-way analysis of variance, followed by a Šidák multiple-comparison test. **f.** Migration speed of tracked nuclei of LGR5- and LGR5+ cells, divided in categories according to the confinement response. Data are represented as the mean ± s.d. of n = 359 cells from four independent experiments. Statistical significance was determined using two-way analysis of variance, followed by a Šidák multiple-comparison test. **g.** Relative mRNA expression levels of *Krt20*, *Lgr5*, and ERM proteins for sorted PDO-p18 cells. Data are represented as the mean ± s.d. of triplicates from one experiment.

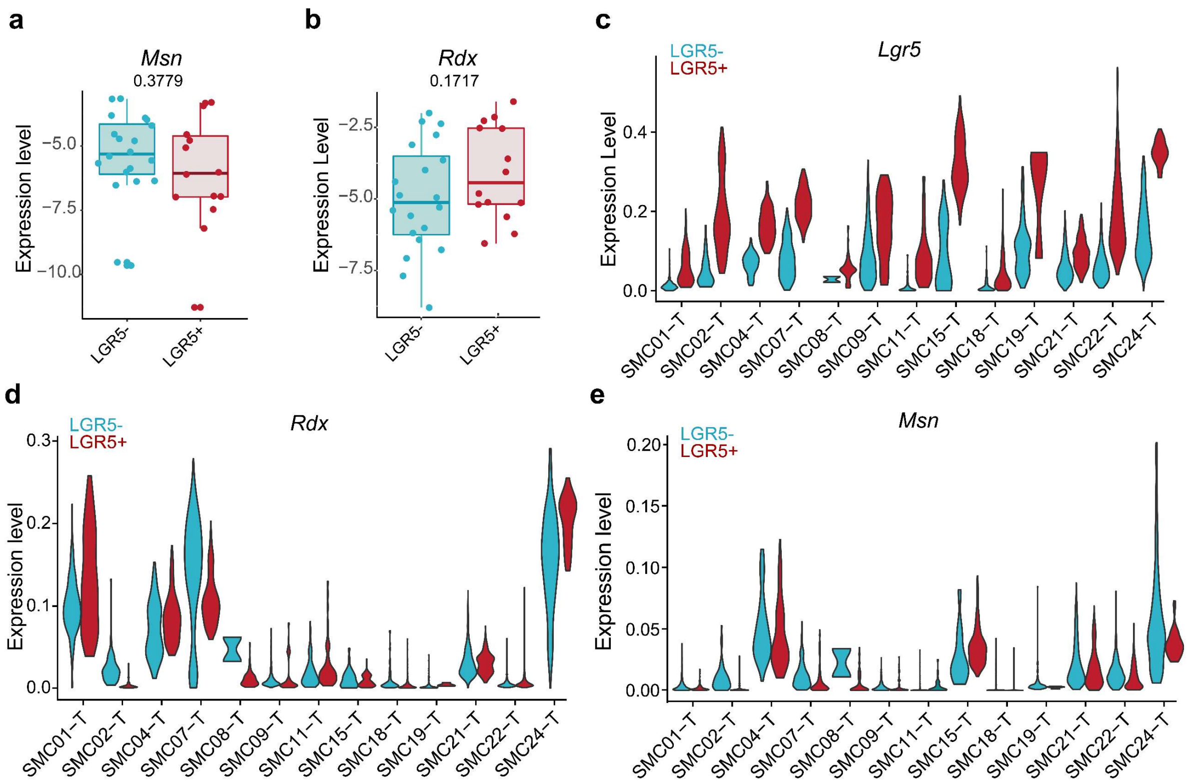

**Supplementary Figure 4**. **Expression of *Lgr5*, *Msn* and *Rdx* in CRC patients.** (**a, b**) Gene expression levels of *Rdx* and *Msn* in epithelial tumor cells from CRC patients in the SMC cohort summarized by patient through the average. Each dot corresponds to the average expression levels of one patient. summarized by patient through the average. The boxes center line represents the median. The box limits represent the first and third quartiles. Whiskers indicate maximum and minimum values. n = 15. (**c-e)** Violin plots showing expression levels of *Lgr5* (**c**), *Rdx* (**d**) and *Msn* (**e**) in epithelial tumor cells from patients in the SMC cohorts. Patient ID is detailed.

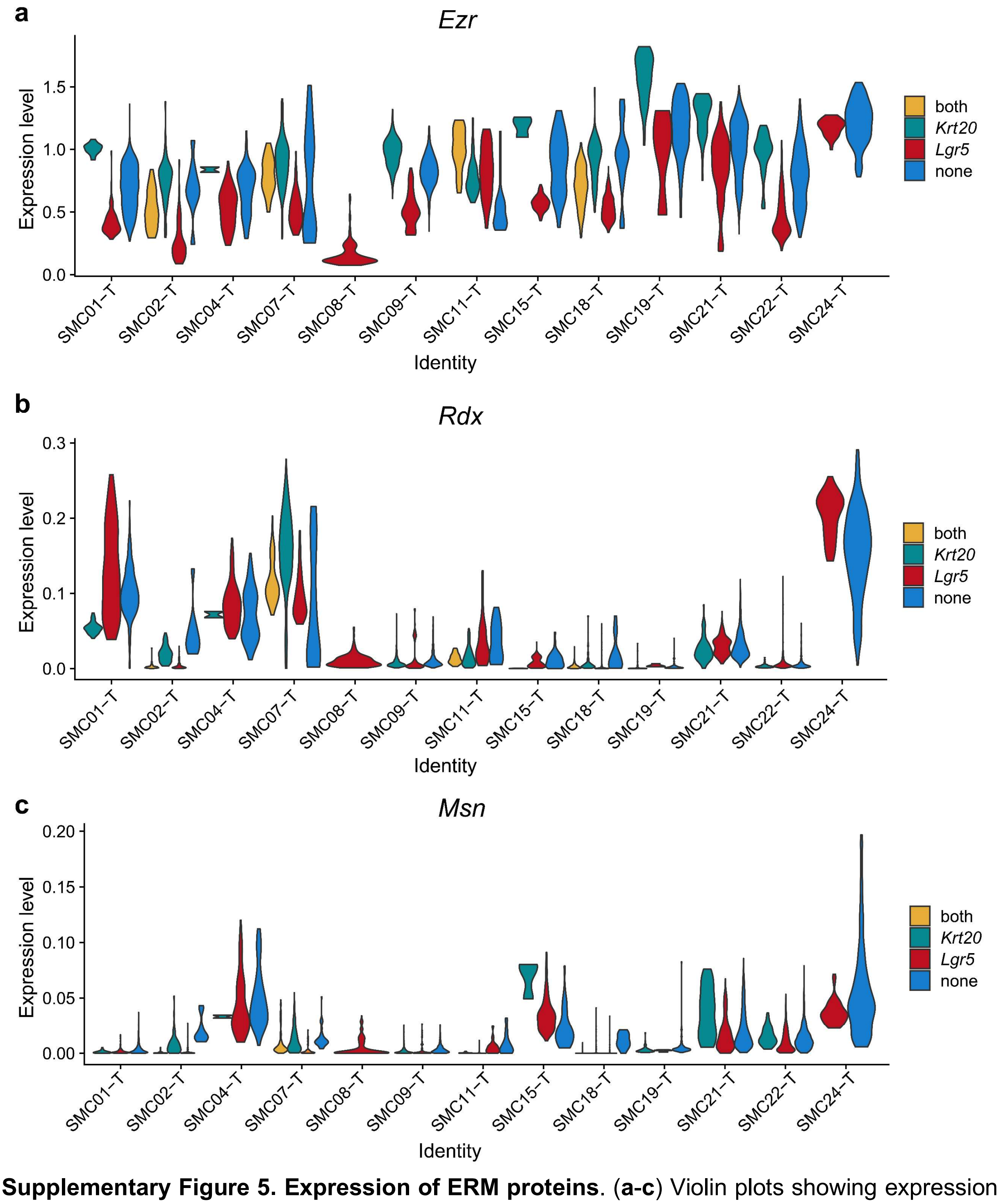

**Supplementary Figure 5. Expression of ERM proteins**. (**a-c**) Violin plots showing expression levels of *Ezr* (**a**), *Rdx* (**b**) and *Msn* (**c**) in epithelial tumor cells from patients in the SMC cohorts. Patient ID is provided in the horizontal axis. Cells were divided in four groups depending on whether they expressed *Lgr5* onl*y (red), Krt20* only (green), both (yellow) or none (blue).

**Supplementary Table**

Table 1.
*p* values for NES analysis.

| **Selected (Supp. Fig. 1c)** | | | |
|---|---|---|---|
| | **LGR5+ vs. LGR5-** | **LGR5+ vs. LGR5$^{med}$** | **LGR5$^{med}$ vs LGR5-** |
| Differentiation (diffmix) | 0.00029 | 0.22100 | 0.00037 |
| GOBLET_Nusse_2018 | 0.00029 | 0.14538 | 0.00037 |
| ENTEROCYTE_Nusse_2018 | 0.00029 | 0.09419 | 0.00037 |
| Mucosecreting_Hs | 0.00029 | 0.00176 | 0.01232 |
| TUFT_Nusse_2018 | 0.00029 | 0.14538 | 0.00037 |
| coreHRC 100 (EpiHR) | 0.00029 | 0.18411 | 0.00037 |
| Fetal signature mustafa | 0.00029 | 0.11702 | 0.00037 |
| Lgr5.ISC.refined | 0.00029 | 0.20591 | 0.02243 |
| POLR1A.High | 0.00029 | 0.14538 | 0.00037 |
| Mex3aHI LGR5hi signature SI | 0.00029 | 0.14538 | 0.03422 |
| Proliferation | 0.00029 | 0.14297 | 0.00037 |
| Ki67_HvL | 0.00029 | 0.10460 | 0.00037 |
| **GOBP gene sets (Supp. Fig. 1d)** | | | |
| substrate adhesion-dependent cell spreading | 0.00041 | 0.33544 | 0.00083 |
| EMT | 0.00041 | 0.23567 | 0.00083 |
| epithelial cell migration | 0.00041 | 0.31564 | 0.13335 |
| cytoskeleton organization | 0.00041 | 0.32075 | 0.05349 |
| PM organization | 0.00041 | 0.10397 | 0.11037 |
| **GOCC gene sets (Fig. 5a)** | | | |
| anchored to membrane | 0.00046 | 0.87661 | 0.00087 |
| cell cortex | 0.00040 | 0.36139 | 0.00087 |
| actin filament bundle | 0.02235 | 0.35013 | 0.05499 |
| actin filament binding | 0.00046 | 0.55087 | 0.11279 |
| PDZ domain binding | 0.00040 | 0.29591 | 0.00089 |
| $PIP_2$ binding | 0.00040 | 0.27463 | 0.20831 |

Table 2.
Primer sequences for RT-qPCR.

| **Primer** | **Sequence** |
|---|---|
| *Lgr5* pair 1 forward | GCTTCCTGGAGGAGTTACGTC |
| *Lgr5* pair 1 reverse | AACAGCTTGGGGGCACATAG |
| *Krt20* pair 6 forward | CAGTGGTACGAAACCAACGC |
| *Krt20* pair 6 reverse | TCCTCTCTCAGTCTCATACTTCAG |

| | |
|---|---|
| *Ezr* pair 1 forward | CTGCTCTGACTCCAGGTTGG |
| *Ezr* pair 1 reverse | GCCGATAGTCTTTACCACCTGAT |
| *Rdx* pair 7 forward | GATGAGTTTGAAGCAATGTGGGG |
| *Rdx* pair 7 reverse | TTAAGGCCCCAGAAAAACCCA |
| *Msn* pair 1 forward | CCATGCCCAAAACGATCAGTG |
| *Msn* pair 1 reverse | CAGCCAGGTGGAGAAACCTT |

Table 3.
shRNA sequences and vectors used to obtain silencing of ERM proteins

| Gene Name | Refseq | CloneID | Target Seq | OligoSeq | vectorID |
|---|---|---|---|---|---|
| VIL2 | NM_003379.3 | TRCN0000062460 | CCCACGTCTGAGAATCAACAA | CCGGCCCACGTCTGAGAATCAACAACTCGAGTTGTTGATTCTCAGACGTGGGTTTTTG | pLKO.1 |
| VIL2 | NM_003379.3 | TRCN0000062461 | CGTGGGATGCTCAAAGATAAT | CCGGCGTGGGATGCTCAAAGATAATCTCGAGATTATCTTTGAGCATCCCACGTTTTTG | pLKO.1 |
| RDX | NM_002906.3 | TRCN0000062435 | GCCAGAGATGAAACCAAGAAA | CCGGGCCAGAGATGAAACCAAGAAACTCGAGTTTCTTGGTTTCATCTCTGGCTTTTTG | pLKO.1 |
| MSN | NM_002444.2 | TRCN0000062411 | GCATTGACGAATTTGAGTCTA | CCGGGCATTGACGAATTTGAGTCTACTCGAGTAGACTCAAATTCGTCAATGCTTTTTG | pLKO.1 |

Table 4.
Different recipes used for the PAA gel preparation.

| Stiffness (kPa) | Acrylamyde (BioRad) % | Bis-acrylamide (BioRad) % | Beads** % solids | Ammonium persulphate (Sigma-Aldrich) % | Tetramethylethylenediamine (Sigma-Aldrich) % |
|---|---|---|---|---|---|
| 0.5 | 4 | 0.03 | 0.03 | 0.5 | 0.05 |
| 1.5 | 5 | 0.04 | 0.03 | 0.5 | 0.05 |
| 3 | 6.16 | 0.044 | 0.03 | 0.5 | 0.05 |
| 5 | 7.46 | 0.044 | 0.03 | 0.5 | 0.05 |
| 11 | 7.5 | 0.1 | 0.03 | 0.5 | 0.05 |
| 30 | 12 | 0.15 | 0.03 | 0.5 | 0.05 |

**Supplementary Movies**

**Supplementary Movie 1. CRC PDOs differentiate *in vitro*.**
Confocal z-stack of a representative PDO cultured for 1 week in culture matrix gel stained for cytokeratin 20 (cyan). LGR5+ cells are labelled with Tdtomato (red).

**Supplementary Movie 2. Response to confinement of LGR5+ and LGR5- cells.**
Representative example of confined PDO single cells on a non-adhesive surface. LGR5+ cells are marked in red while LGR5- cells are unlabeled. All nuclei are labeled (blue). Images were acquired every 5 minutes for a total duration of 100 min.

**Supplementary Movie 3. Response to confinement of LGR5+ and LGR5- cells.**
Second representative example of confined PDO single cells on a non-adhesive surface. LGR5+ cells are marked in red while LGR5- cells are unlabeled. All nuclei are labelled (blue). Images were acquired every 5 minutes for a total duration of 100 min.

**Supplementary Movie 4. Tracking of polarized LGR5- and LGR5+ cells.**
Representative experiment . Left: LGR5+ cells are marked in red while LGR5- cells are unlabeled. All nuclei are labelled (blue). Right: cellular tracks overlaid on the phase contrast images. Images were acquired every 5 minutes for a total duration of 100 min. Note the fast LGR5- cell with a blue track.

**Supplementary Movie 5. Tracking of polarized LGR5- and LGR5+ cells.**
Second representative experiment. Left: LGR5+ cells are marked in red while LGR5- cells are unlabeled. All nuclei are labelled (blue). Right: cellular tracks overlaid on the phase contrast images. Images were acquired every 5 minutes for a total duration of 100 min.

**Supplementary Movie 6. $LGR5^{high}$ cluster migrating on 2D soft substrate.**
Left: Time lapse of $LGR5^{high}$ PDO cluster was acquired every hour for 15 hours. LGR5+ cells are marked in red while LGR5- cells are unlabeled. Right: Time lapse of cellular tractions corresponding to the cluster shown on the left panel. Colormap and traction vector scale are the same as in Fig. 3B.

**Supplementary Movie 7. $LGR5^{low}$ cluster migrating on 2D soft substrate.**
Left: Time lapse of $LGR5^{low}$ PDO cluster was acquired every hour for 15 hours. LGR5+ cells are marked in red while LGR5- cells are unlabeled. Right: Time lapse of cellular tractions corresponding to the cluster shown on the left panel. Colormap and traction vector scale are the same as in Fig. 3B.

**Supplementary Movie 8. $LGR5^{low}$ cluster attaching to an endothelial monolayer.**
Time lapse of $LGR5^{low}$ PDO cluster seeded on top of a HUVEC monolayer. LGR5+ cells are marked in red and HUVEC cells are marked in green. Images were acquired 1 h after cluster seeding, every 40 minutes for 14 h.

**Supplementary Movie 9. $LGR5^{med}$ cluster attaching to an endothelial monolayer.**
Time lapse of $LGR5^{med}$ PDO cluster seeded on top of a HUVEC monolayer. LGR5+ cells are marked in red and HUVEC cells are marked in green. Images were acquired 1 h after cluster seeding, every 40 minutes for 14 h.

**Supplementary Movie 10. $LGR5^{high}$ cluster attaching to an endothelial monolayer.**

Time lapse of LGR5$^{high}$ PDO cluster seeded on top of a HUVEC monolayer. LGR5+ cells are marked in red and HUVEC cells are marked in green. Images were acquired 1 h after cluster seeding, every 40 minutes for 14 h.

**Supplementary Movie 11. Response to confinement of LGR5+ cells expressing the synthetic iMC-linker and LGR5+ cells.**
Confined PDO single cells on a non-adhesive surface. LGR5+ cells are marked in red. All nuclei are labelled (blue). Images were acquired every 5 minutes for a total duration of 70 min.
